# Regenerative Agriculture Augments Bacterial Community Structure for a Healthier Soil and Agriculture

**DOI:** 10.1101/2022.11.06.515329

**Authors:** Indira Singh, Meeran Hussain, Manjunath G, Nagasuma Chandra, Ravikanth G

**Author notes:** For Correspondence, Indira Singh: Senior Development Officer, Ashoka Trust for Research in Ecology and the Environment (ATREE), Royal Enclave, Srirampura, Jakkur, Bangalore, Karnataka – 560064, India.

## Abstract

Use of chemical fertilization and pesticides not only harm the environment but also have detrimental consequences on human health. In recent years, there has been a major emphasis worldwide on natural agriculture methods. Regenerative agriculture is known across the world as a combination of nature-friendly farming practices such as no-till, cover cropping, crop-rotation, agro-forestry and use of organic home-based/farm-based ingredients to revive soil health. In India, a number of farmers are slowly adopting these practices using home-based mixtures and farmyard manure for soil rejuvenation and pest management. In order to evaluate the efficacy of the regenerative agriculture practices, this study compared conventional and regenerative agriculture plots for their soil bacterial and nutrient profiles. Two crops - ragi and vegetable (tomato/beans), and different lengths (≤3 and >5 years) of regenerative practices were additional metrics considered to understand variabilities due to crop-type and period of application. We found that all regenerative practices were effective in bringing about an enrichment for soil bacteria with a more heterogeneous composition. Additionally, the regenerative vegetable (RV) plots had an enhanced representation of *Actinobacteriota, Chloroflexi, Cyanobacteria* and *Patescibacteria* in comparison to conventional vegetable (CV) plots and Barren land (BL). Similarly, the regenerative ragi (RR) plots saw higher representation of *Firmicutes* and *Actinobacteriota* in comparison to conventional ragi (CR) plots and BL. The RV plots were also found to be enriched for Plant Growth Promoting Rhizobacteria (PGPRs) - *Pseudomonas sp*., and RR plots were enriched for *Bacillus sp*., and *Mesorhizobium sp*., which are known to play significant roles in vegetable and ragi growth respectively. Interestingly, long-term regenerative agriculture was able to support good nutrient composition while enhancing Soil Organic Carbon (SOC) levels. In all, the regenerative agriculture practices were found to be effective in improving bacterial community structure and simultaneously improving soil health. We found that BL soil with eucalyptus plantation showed least bacterial diversity suggesting detrimental impact on soil health.

## Introduction

Agriculture is the primary livelihood means for more than 50% of India’s population (1). With the advent of green revolution, farmers used conventional agriculture involving intensive use of synthetic fertilizers and pesticides for crop and field management (2, 5). Conventional agriculture with other unsustainable land management practices such as tilling, leaving the soil barren during non-growing season, agricultural intensification and monoculture cropping have led to the deterioration of soil quality and crop health, leaving the farmers economically distressed (2, 3).

However, there is little scientific evidence regarding the regenerative agricultural practices and their ability to improve soil and crop health. A healthy soil is supported by a robust and thriving microbial community, which can carry out a host of biogeochemical activities to enrich the soil with essential nutrients and plant growth promoters (4, 5, 82). In this study, we compare two farming systems (regenerative and conventional) based on their soil nutrient and bacterial profiles to verify their abilities in restoring soil health in the context of Karnataka’s semi-arid farmlands.

Conventional agriculture, which involves application of chemical fertilizers (Nitrogen, Phosphorus and Potassium, NPK) for boosting agricultural outputs, has been implicated for acidification and deterioration of soil and climate change (6). Excessive addition of nitrogen fertilizer brings about leaching of nitrogen into waterbodies, a major cause of eutrophication apart from accumulation and release of nitrous oxide from soil, a potent greenhouse gas. In contrast, regenerative agriculture uses environment friendly soil and crop management systems, which has the ability to heal the environment cost effectively with minimal inputs (7, 8, 9, 10). This soil management technique uses a combination of methods such as no-till, cover cropping, crop rotation, multi and inter-cropping, mulching and farm-based manure application. Overall, regenerative agriculture uses only naturally available inputs for improving soil health and is proposed to help in mitigating climate change by enhancing the soil’s carbon storage capacity (9, 10).

Some of India’s smallholder farmers have recently started to adopt regenerative agriculture to improve their soil and crop health. Alongside using the globally practiced regenerative methods, smallholders in Karnataka also use soil-rejuvenation methods based on traditional knowledge. Homemade additives made from cow-products and other easily available ingredients such as jaggery and chickpea flour. Although, there is a huge repertoire of knowledge accumulating to show the benefits of regenerative agricultural system, yet there is an ongoing debate on integrating the two systems to achieve sustainability in food production (7, 9). Consistent with this idea, many Indian farmers use both chemical fertilizers and farm-based manure for better yield (11, 12). This study attempts to assess the impact of merging the two systems on the soil’s bacterial composition.

The soil microbial community is comprised of bacteria, fungi, viruses and protozoans. These microbes carry out the fundamental processes facilitating-nutrient cycling, decomposition of organic matter, defining soil texture, soil water-retention capacity, degradation of toxic wastes and preventing the growth of plant pests and pathogens (13). Different soil treatments can have an impact on the microbial community structure, but the microbiome changes are very complex processes stimulated by multiple factors such as temperature, climate, additives/treatments, type of crop grown, cropping patterns, etc. Sustainable agriculture practices should ideally boost the growth and prevalence of beneficial microbes over the pathogenic species. Studies show that regenerative agriculture manifests soil health by improving soil microbial diversity and richness (14, 15, 16, 17). However, availability of too many regenerative agriculture options with little knowledge about their anticipated outcomes, followed by a long time-period for a demonstrable change in soil health/plant yield, makes a smallholder farmer desperate and vulnerable. Therefore, a scientific understanding of the basis of soil health promotion by these practices is essential for enabling an evidence-based recommendation. Additionally, due to availability of a broad range of regenerative practices, along with huge variabilities in regional soil types, climatic conditions, timing and extent of application and differences in crop type and cropping patterns, it is extremely difficult to compare studies from across the world. Therefore, a region specific and country specific study would be useful to obtain first-hand information on the mode of action and benefits accrued. To date there is no such study reported from India to show the comparative advantage of using regenerative agriculture on soil microbial diversity.

Metagenomics analysis using Next Generation High-throughput sequencing of soil DNA samples has been an efficient tool to determine the microbiome in soil. The technique provides details on the diversity, abundance and occurrence of specific genera and species in the given sample (15, 18, 19, 20, 21). Here, using 16S metagenomics, we compared the bacterial community structure under regenerative agriculture with that observed in conventional agriculture and barren land. Further, the metagenomics datasets were analyzed for alpha and beta diversity to establish the bacterial diversity in different samples. Agricultural plots growing either vegetable crops (tomato/bean) or finger-millet crop (Ragi) were considered for this study.

We found that agriculture plots following regenerative methods recorded an enhancement in bacterial diversity, enriched for specific plant growth promoting bacterial genera compared to conventional agriculture plots and barren land. The results from this study provide strong evidence to show the significance of regenerative agriculture in boosting soil microbial health to improve healthy nutrient composition, organic carbon content, water retention property and consequently induce plant growth and productivity. Our study indicates that long term and regular use of regenerative farm practices by farmers in Karnataka will have potential to support sustainability in soil health and agriculture.

## Materials and Methods

### Soil Sample Collection

This study aimed to establish the impact of regenerative agriculture practices on soil nutrient composition and microbial health with respect to the number of years of application. We considered two types of crops for this study – Ragi (finger-millet) and Tomato/Bean (Vegetable) crop. Soil sampling was carried out in January and February of 2021 when there was a brief respite from Covid-19. Therefore, some samples were collected in absence of the crop. Soil was collected from near the roots of the crops wherever we could find plots with crops and for others soil was collected at the depth of 1-5 cm from the top. We collected soil from four corners of the plots and one from the center of the plot. Finally, all the soil samples from one plot were pooled together for experimentation. For physicochemical analysis, we collected about 2 kg of the soil pooling soil samples from all the five locations on the plot into one common bag. For the microbiome study soil was collected in sterile falcon tubes kept on ice and finally stored at −20 ^0^C until further processing. Soil sampling was done as given in *Table 1*.

**Table 1.**
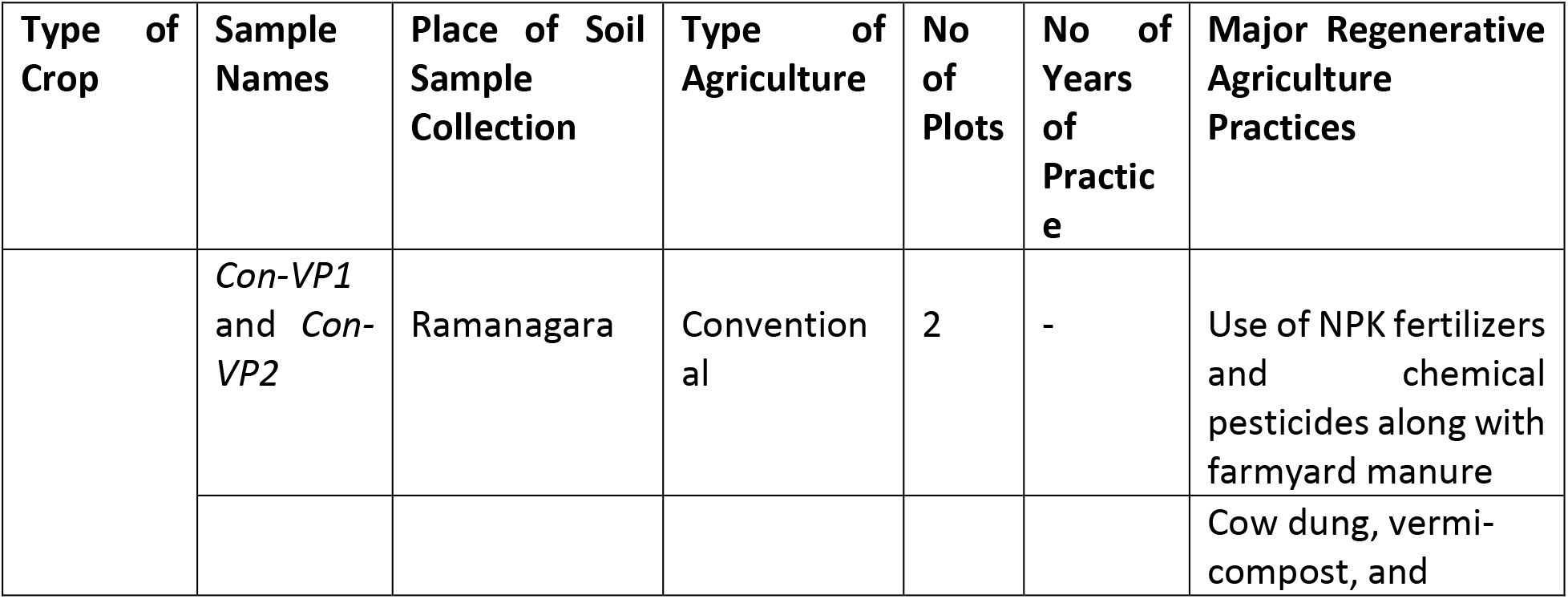
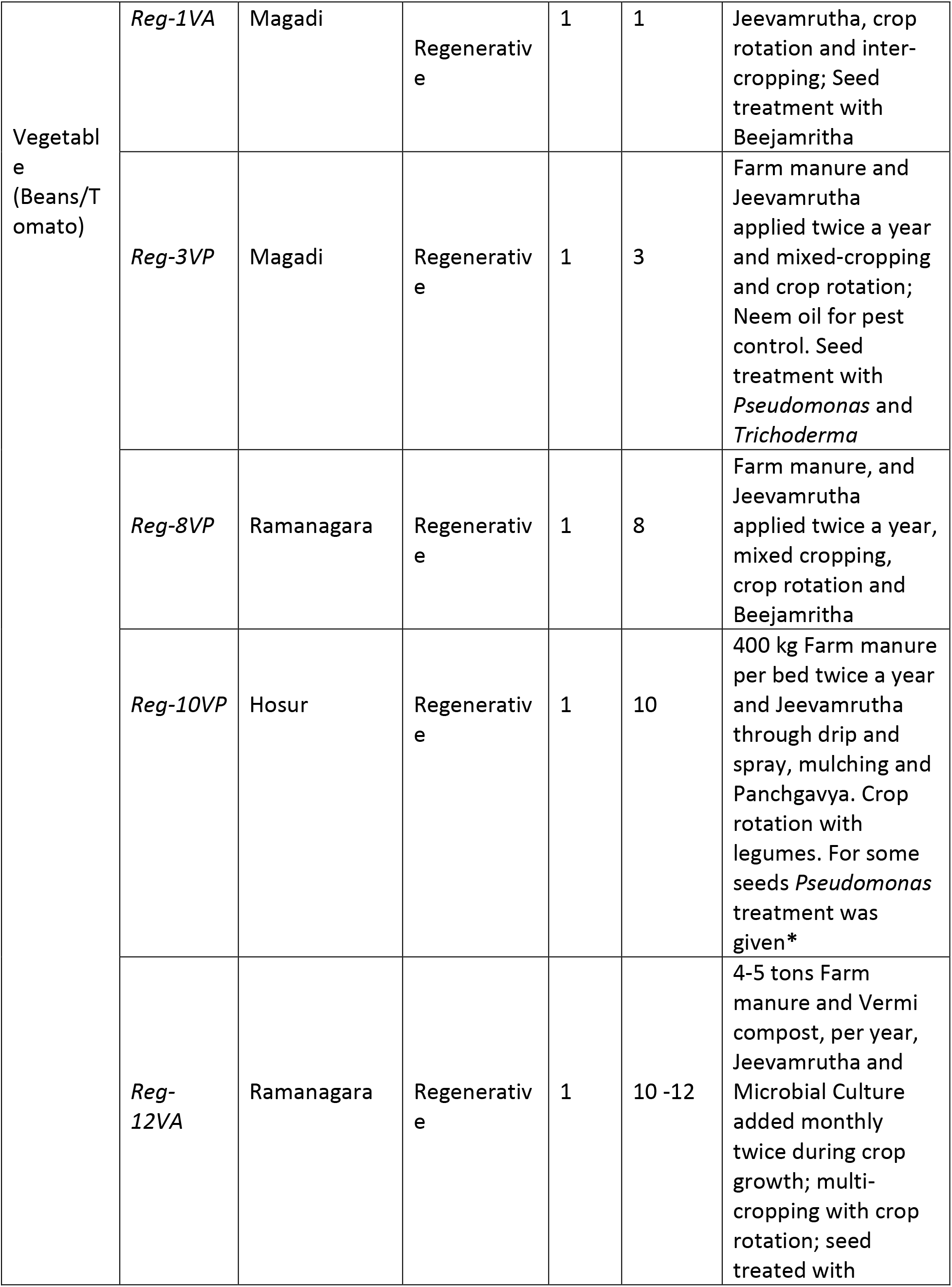
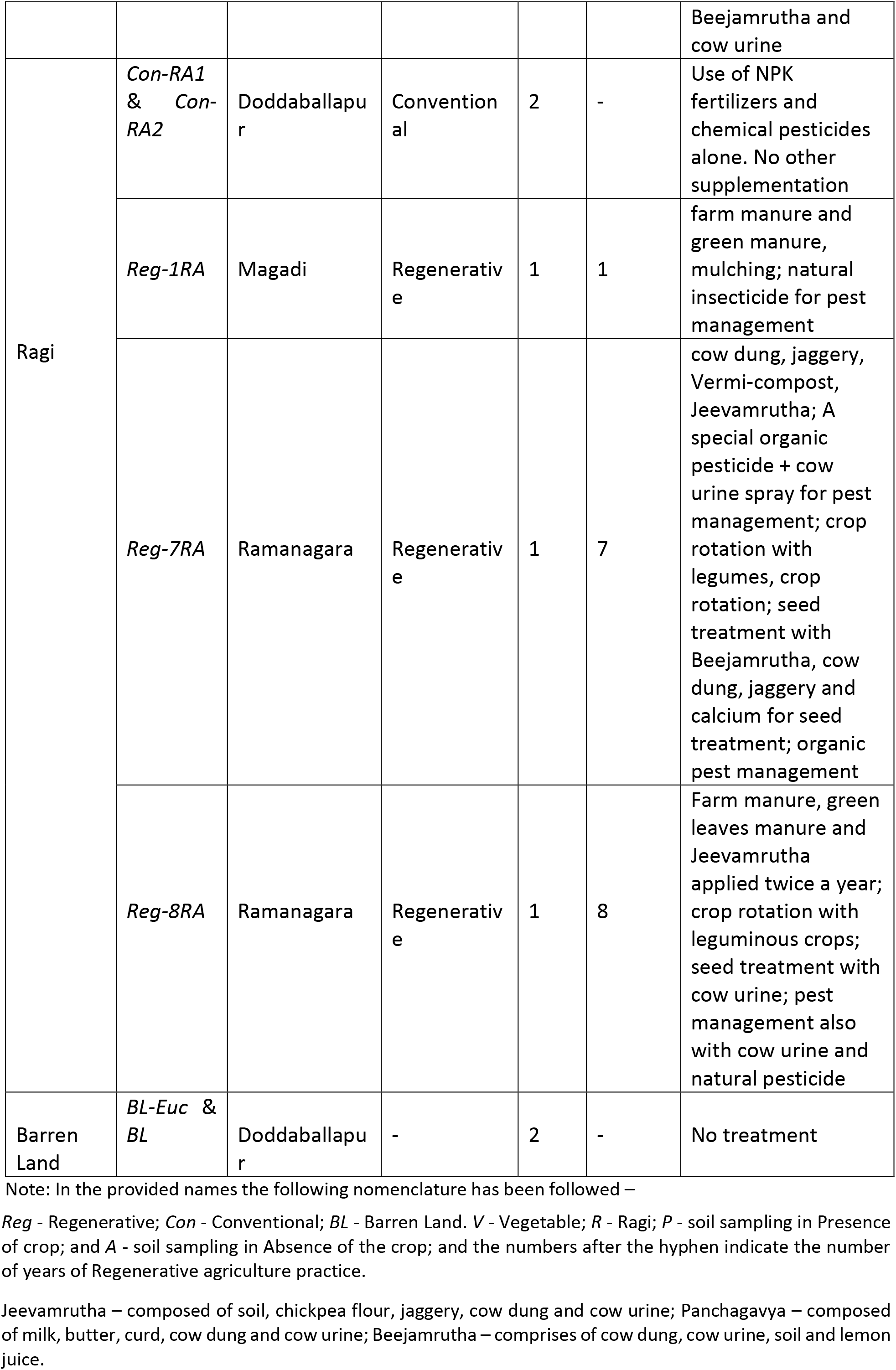
Soil Sampling.

We selected the plots for this study in the outskirts of Bengaluru in the towns of Ramanagara, Magadi, Doddaballapur and Hosur. This region is predominantly semi-arid. Barren land (BL) samples with no vegetation and with eucalyptus formed the no treatment controls. Barren land with eucalyptus (*BL-Euc*) was included as an additional metric in the study to get a sense of how monocultures impact soil health. The regenerative plots varied greatly in the kind of application practiced. For instance, some farmers used farmyard manure and Jeevamrutha, while others used farmyard manure, Jeevamrutha along with vermicompost (*Table 2*).

**Table 2.** Physicochemical Parameters of the Soil Samples.

| Sample Name | pH | EC (dS/m) | Organic carbon (%) | Nitrogen | Phosphorus | Potassium | Calcium | Magnesium | Zinc | Manganese | Iron | Copper |
| --- | --- | --- | --- | --- | --- | --- | --- | --- | --- | --- | --- | --- |
|  |  |  |  | kg/ha |  |  | mEq/1000 g |  | ppm |  |  |  |
| IDEAL | 6.5-7.5 | <1.00 | 0.5-0.75 | 280-560 | 22.9-56.33 | 141-336 | >1.5 | >1.0 | >0.6 | >2.0 | 2.5 - 4.5 | >0.2 |
| *Con-VP1* | 7.54 | 0.367 | 0.39 | 131.8 | 342.47 | 250.3 | 38 | 21 | 3.72 | 7.2 | 35.64 | 1.17 |
| *Con-VP2* | 7.6 | 0.399 | 0.44 | 106.6 | 346.28 | 284.4 | 41 | 27 | 4.77 | 7.68 | 11.31 | 0.84 |
| *Reg-1VA* | 7.41 | 0.113 | 0.29 | 125.4 | 87.64 | 217.9 | 35 | 19 | 1.23 | 6.33 | 13.32 | 0.42 |
| *Reg-3VP* | 7.43 | 0.193 | 0.3 | 156.8 | 62.39 | 184 | 40 | 30 | 4.44 | 3.99 | 32.16 | 1.56 |
| *Reg-8VP* | 8.31 | 0.267 | 0.36 | 120.1 | 152.8 | 189.7 | 62 | 47 | 2.34 | 6.27 | 8.79 | 0.72 |
| *Reg-10VP* | 7.95 | 0.279 | 0.51 | 144.2 | 510.13 | 506.4 | 105 | 64 | 4.08 | 8.34 | 19.62 | 2.19 |
| *Reg-12VA* | 7.7<br>1 | 0.231 | 0.32 | 100.3 | 187.3<br>3 | 223.5 | 79 | 54 | 2.79 | 9.42 | 16.53 | 0.81 |
| *Con-RA1* | 5.7<br>9 | 0.316 | 0.35 | 131 | 37.6 | 334 | 14 | 6 | 1.32 | 10.8 | 16.56 | 0.45 |
| *Con-RA2* | 3.9<br>4 | 0.159 | 0.39 | 144 | 44.7 | 170 | 21 | 10 | 1.14 | 18.63 | 78.27 | 1.5 |
| *Reg-1RA* | 7.3<br>5 | 0.128 | 0.38 | 106.6 | 58.58 | 216.6 | 58 | 32 | 1.05 | 11.01 | 10.32 | 0.51 |
| *Reg-7RA* | 6.8<br>9 | 0.09 | 0.36 | 119.1 | 148.6<br>1 | 252.7 | 45 | 28 | 2.58 | 14.58 | 36.18 | 1.56 |
| *Reg-8RA* | 7.0<br>1 | 0.13 | 0.42 | 125.4 | 60.01 | 306.7 | 54 | 38 | 2.37 | 16.02 | 23.13 | 0.57 |
| *BL- Euc* | 6.0<br>4 | 0.235 | 0.31 | 119 | 29 | 242 | 27 | 14 | 1.51 | 23.91 | 9.3 | 0.459 |
| *BL* | 5.8<br>5 | 0.106 | 0.41 | 150 | 14.7 | 108 | 22 | 9 | 0.99 | 6.33 | 16.29 | 0.327 |

Note: The ideal values are based on recommendations given by the Indian Society of Soil Science (31).

Sample grouping into categories for analysis:

1. two conventional vegetable (CV) plots – *Con-VP1* & *Con-VP2*
2. two conventional ragi (CR) plots – *Con-RA1* & *Con-RA2*
3. two regenerative (≤3 years) vegetable (RV) plots – *Reg-1VA* & *Reg-3VP*
4. three regenerative (>5 years) vegetable (RV) plots – *Reg-8VP, Reg-10VP* & *Reg-12VA*
5. one regenerative (≤3 years) ragi (RR) plot– *Reg-1RA*
6. two regenerative (> 5 years) ragi (RR) plots – *Reg-7RA* & *Reg-8RA*
7. two barren land samples – *BL* (no vegetation) & *BL-Euc* (with Eucalyptus)

### Soil Physicochemical Analysis

Collected soil samples were taken to the laboratory, shade dried, pounded using wooden pestle and mortar, sieved (2 mm) and stored in airtight polyethylene bags for further analysis. The soil samples were analysed for various electrochemical properties. The soil pH, electrical conductivity, organic carbon content, nutrients namely - nitrogen, phosphorus, potassium, calcium, magnesium, sulphur and micronutrients - iron, zinc, manganese and copper were analyzed according to the standard procedures as given in *Table 3*.

**Table 3.** Methods adopted for soil analysis.

| Sl. No. | Parameter | Method |
| --- | --- | --- |
| 1. | Soil reaction (pH)<br>(1:2.5 soil: water suspension) | Potentiometry (22) |
| 2. | Electrical conductivity<br>(1:2.5 soil: water suspension) | Conductometry (22) |
| 3. | Organic carbon (%) | Wet oxidation (23) |
| 4. | Available Nitrogen (kg ha<sup>-1</sup>) | Macro kjeldahl<br>Distillation (24) |
| 5. | Available Phosphorus (kg ha<sup>-1</sup>) | Spectrophotometry (25) |
| 6. | Available Potassium (kg ha<sup>-1</sup>) | Flame photometry (22) |
| 7. | Exchangeable Calcium and<br>Magnesium (mEq/1000 g) | Complexometric titration (22) |
| 8. | Available Sulphur (ppm) | Turbidometry (22) |
| 9. | DTPA extractable Iron,<br>Manganese, Zinc and Copper<br>(ppm) | Atomic Absorption<br>Spectrophotometry (26) |

### Soil DNA Isolation, library preparation and deep sequencing

DNA was isolated from the soil samples using DNeasy Power soil kit, following manufacturer’s protocol. DNA samples were sent for 16S metagenomics analysis to Eurofins, where amplicon sequencing was done using Illumina MiSeq platform (Eurofins Genomics India Pvt. Ltd., Bangalore, India). The quality of the DNA samples was checked using NanoDrop estimation by determining A260/280 ratio. The amplicon libraries were prepared using Nextera XT Index Kit (Illumina inc.) as per the 16S Metagenomic Sequencing Library preparation protocol (Part # 15044223 Rev. B). Primers for the amplification of the bacterial 16S V3-V4 region were designed and synthesized at Eurofins Genomics Lab. Amplification of the 16S gene was carried out. The QC passed amplicons with the Illumina adaptor were amplified using i5 and i7 primers that add multiplexing index sequences as well as common adapters required for cluster generation (P5 and P7) as per the standard Illumina protocol. The amplicon libraries were purified by AMPure XP beads and quantified using Qubit Fluorometer. The amplified and AMPure XP bead purified libraries were analyzed on 4200 Tape Station system (Agilent Technologies) using D1000 Screen tape as per manufacturer’s instructions. After obtaining the mean peak sizes from Tape Station profile, libraries were loaded onto MiSeq at appropriate concentration (10-20 pM) for cluster generation and sequencing. Paired-end sequencing allows the template fragments to be sequenced in both the forward and reverse directions on MiSeq. Kit reagents were used for binding the samples to complementary adapter oligoes on paired-end flow cell. The adapters were designed to allow selective cleavage of the forward strands after re-synthesis of the reverse strand during sequencing. The copied reverse strand was then used to sequence from the opposite end of the fragment.

### Metagenomics Analysis

In all, there were 14 samples and the number of read pairs ranged from 100,468 to 341,993 per sample. Quality check of 16s rRNA sequences was done using FastQC (v0.11.5) and the adapter sequences were removed using Trimgalore (version: 0.6.7) (27, 28). The complete metagenome analysis was done using the QIIME 2.0 (Quantitative Insights in to Microbial Ecology) (version: 2021.4.0) pipeline (29). De-noising of the paired-end reads was done using the DADA2 tool that is within QIIME 2.0 which is used to filter low-quality reads of Phred score <15. High-quality reads were retained in 16S rRNA sequences by truncating the length of the forward read to 285 bp and the reverse reads to 250 bp. The resulting reads were de-noised to obtain unique sequence variants. DADA2 (version: 2021.4.0) produces “operational taxonomic units (OTUs)” by grouping unique sequences; these are 100% equivalent to the OTUs and are referred to as “Amplicon Sequence Variants (ASVs)”. The feature table was constructed using QIIME 2.0, which is similar to the BIOM table and the representative sequence file.

Further, the phylogenetic tree was built for each sample using the MAFT program, which is an inbuilt plugin in the QIIME 2.0 pipeline, results from this program are used to study the Alpha diversity by using Faith’s Phylogenetic and Pielou’s evenness matrix. Alpha diversity is further explored as a function of sampling depth by performing Alpha Rarefaction. Taxonomic classification was done by mapping the sequences at 99% sequence identity to an optimized version of the SILVA database using Naive Bayes classifier and q2-feature-classifier plugin of QIIME 2.0. The results of each step were downloaded from the QIIME2 program and they were plotted using ggplot2 (3.3.5) with R programming language (29).

## RESULTS

### Soil’s organic carbon and major nutrient composition

The physicochemical properties of soil such as - pH, major and minor nutrient composition obtained in the study were compared with pre-defined ideal values (given in Table 3). The results from the soil physicochemical analysis show that except *Con-RA2* (pH = 3.94), *Con-RA1* (pH = 5.79), *BL* (pH = 5.85) and *BL-Euc* (pH = 6.04), all other samples had pH either in the ideal range (6.5-7.5) or in the moderately alkaline range (31).

For most parameters, there was no significant difference between the conventional and regenerative agriculture plots. For instance, nitrogen levels were observed to be much less than the required range of 280 – 560 kg/ha in all the plots. Phosphorus levels were much above the required range of 22.9 – 56.3 kg/ha, while potassium was in the ideal range (141-3663 kg/ha) in all the soil samples. An important finding was that phosphorus and potassium are present at very high levels in *Reg-10VP* soil with the use of only organic manure. The *Reg-10VP* plot uses very heavy application of cattle manure and other household+ farm-based mixture and has been using these practices for as long as 10 years. It would be interesting to study how cattle manure and each of these practices individually contribute to soil’s phosphorus and potassium content. Additionally, *Reg-10VP* also showed the best organic carbon composition of 0.51% (ideal – 0.5 – 0.75%), unlike all other soil samples which remained below the ideal range. In contrast, the other regenerative agriculture plots in this study did not seem to show such a remarkable enhancement in their nutrient profiles when compared with the conventional agriculture soil. However, most regenerative plots have desired levels of most macro- and micronutrients barring nitrogen and organic carbon levels. This clearly indicates that most of these regenerative soil treatments regimens have the ability to provision maximum of these nutrients even in the absence of inorganic additives.

Further investigations will be needed to establish the basis for the improved physicochemical profiles in *Reg-10VP* soil. Altogether, these findings suggest that the long-term application of regenerative practices could help to improve the soil’s nutrient composition including organic carbon levels.

### Taxonomic composition of soil microbial community

To identify the bacterial community structure associated with conventional versus regenerative practices, we performed 16S metagenomics studies. A total of 2,941,473 raw sequence reads from 14, 16S metagenome libraries were generated by the Illumina platform, ranging from 1,51,169 to 3,41,993 reads per sample. After removal of adapter sequences, ambiguous reads (reads with unknown nucleotides “N” larger than 5%), and low-quality sequences (reads with QV <20 phred score) and a minimum length of 100 bp, 2,801,991 high quality clean reads were further used for analysis.

The datasets were analyzed with QIIME 2.0 pipeline, using the SILVA database. At phylum level, Proteobacteria, Bacteroidota, Planctomycetota, Cyanobacteria, Actinobacteriota, Chloroflexi, Acidobacteriota, Verrucomicrobiota, Firmicutes and Gemmatimonadetes are the top 10 predominant phyla.

### Bacterial richness and community heterogeneity

Soil samples were classified into following groups for this analysis –

i. Barren (comprising *BL* and *BL-Euc*);
ii. Conv (Vegetable plots - *Con-VP1* and *Con-VP2*) and (Ragi plots - *Con-RA1* and Con-RA2);
iii. Reg≤3 (Vegetable plots - *Reg-1VA* and *Reg-3VP*) and (Ragi plots – *Reg-1RA*);
iv. Reg>5 (Vegetable plots – *Reg-8VP, Reg-10VP* and *Reg-12VA*) and (Ragi plots – *Reg-7RA* and *Reg-8RA*)

For both crop types (vegetable and ragi), we found that regenerative agriculture plots in general showed higher bacterial richness compared to conventional and barren (*Figure 1a*, *1c*). Furthermore, bacterial species evenness comparison showed that both regenerative vegetable (RV) and regenerative ragi (RR) plots displayed least species evenness implying that the species composition in these plots is highly heterogeneous (*Figure 1b*, *1d*). Surprisingly, CR plots showed least bacterial richness (*Figure 1c*) which was even less than that observed in the BL soil, whereas CV soil demonstrated better bacterial richness than BL samples (*Figure 1a*). On a similar note, CR plots had the highest species evenness followed by BL plots (*Figure 1d*), while CV plots had lower species evenness than BL (*Figure 1b*). Our findings indicate that regenerative agriculture increases soil’s bacterial richness and heterogeneity irrespective of crop type and the kind of regenerative practices adopted.

**Figure 1.**
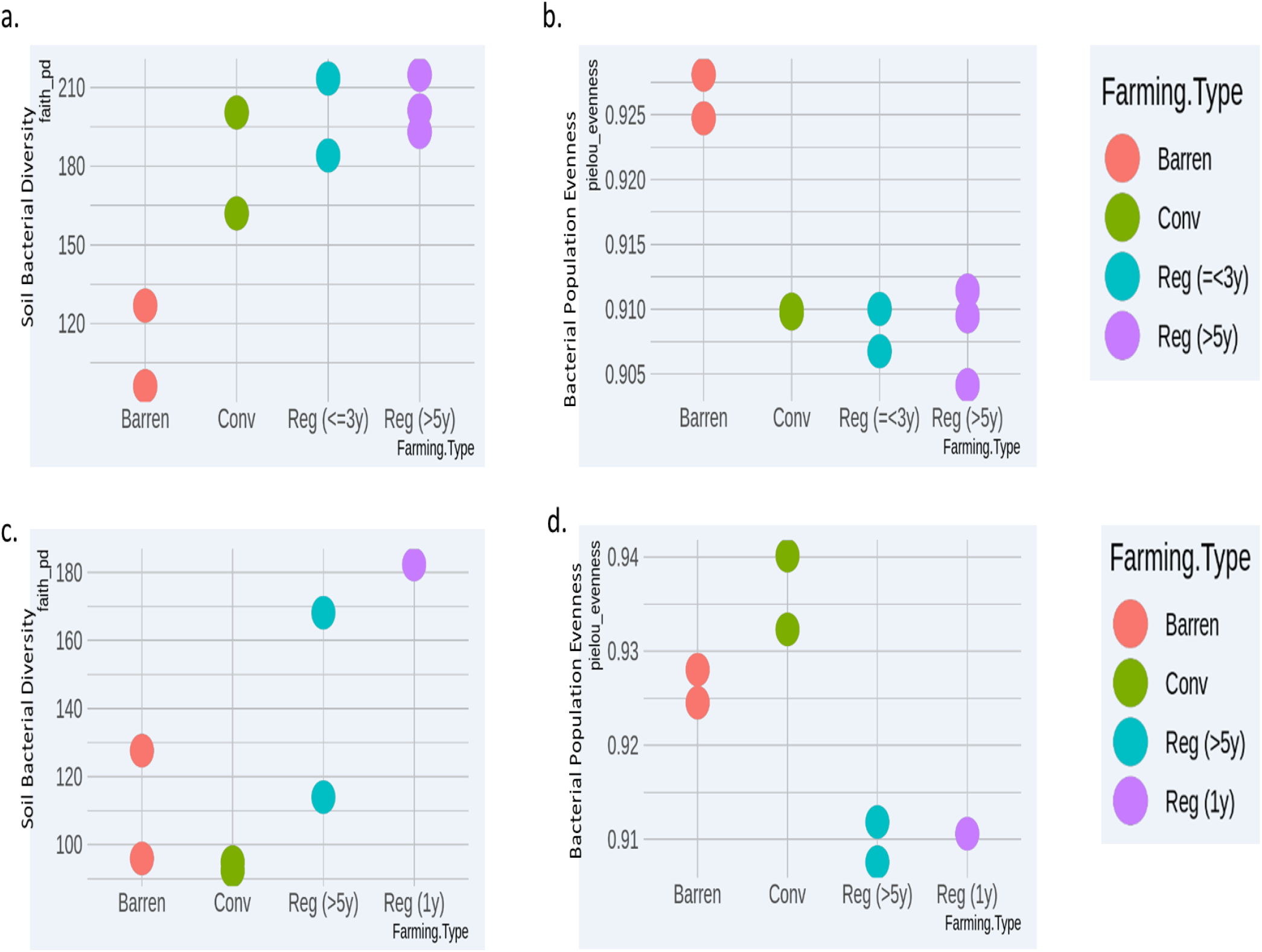
Comparative bacterial Richness **(a)** and Evenness **(b)** analysis of Vegetable growing conventional (*Conv*) and Regenerative agriculture (*Reg*) plots with Barren land (BL) soil. Comparative bacterial Richness **(c)** and Evenness **(d)** analysis of Ragi growing conventional (*Conv*) and Regenerative agriculture (*Reg*) plots with Barren land (BL) soil.

### Alpha diversity

The alpha diversity among different soil samples was compared to determine the mean species diversity in each plot. A higher alpha diversity value therefore signifies a more diverse pool of bacterial species accumulation. It is important to point out here that we collected a few soil samples from regenerative plots in the presence of vegetable crops labeled with the suffix *VP*, in the presence of ragi are labeled as *RP* and those taken post-harvest are labeled with the suffix V*A and RA* respectively. While all CV plot soils were collected in the presence of the crop, all CR plot soils were collected in the absence of the crop.

Overall, the alpha diversity study showed that most regenerative agriculture plots demonstrated higher alpha diversity compared to conventional agriculture plots and barren soil (*Figure 2a, 2b*). Among vegetable plots our results indicate that alpha diversity is directly proportional to the length of regenerative agricultural practice. For example, the bacterial diversity in soil from vegetable regenerative plot practicing for 10 years (*Reg-10VP*) was greater than that observed for the plot practicing for 8 years (*Reg-8VP*) (*Figure 2A*). Likewise, among the post-harvest category, we observed greater bacterial diversity in *Reg-12VA* (12 years) as compared to *Reg-1VA* (1 year) (*Figure 2a*). Surprisingly, and in contrast to time-dependency, *Reg-3VP* (3 years) showed a better alpha diversity than *Reg-8VP* (8 years). We believe that this variability is due to the inherent differences in the soil quality associated with various locations. As expected, soil collected from RA plots where vegetable crops were present showed greater diversity than RA soil samples collected post-crop harvest (*Figure 2a*).

**Figure 2.**
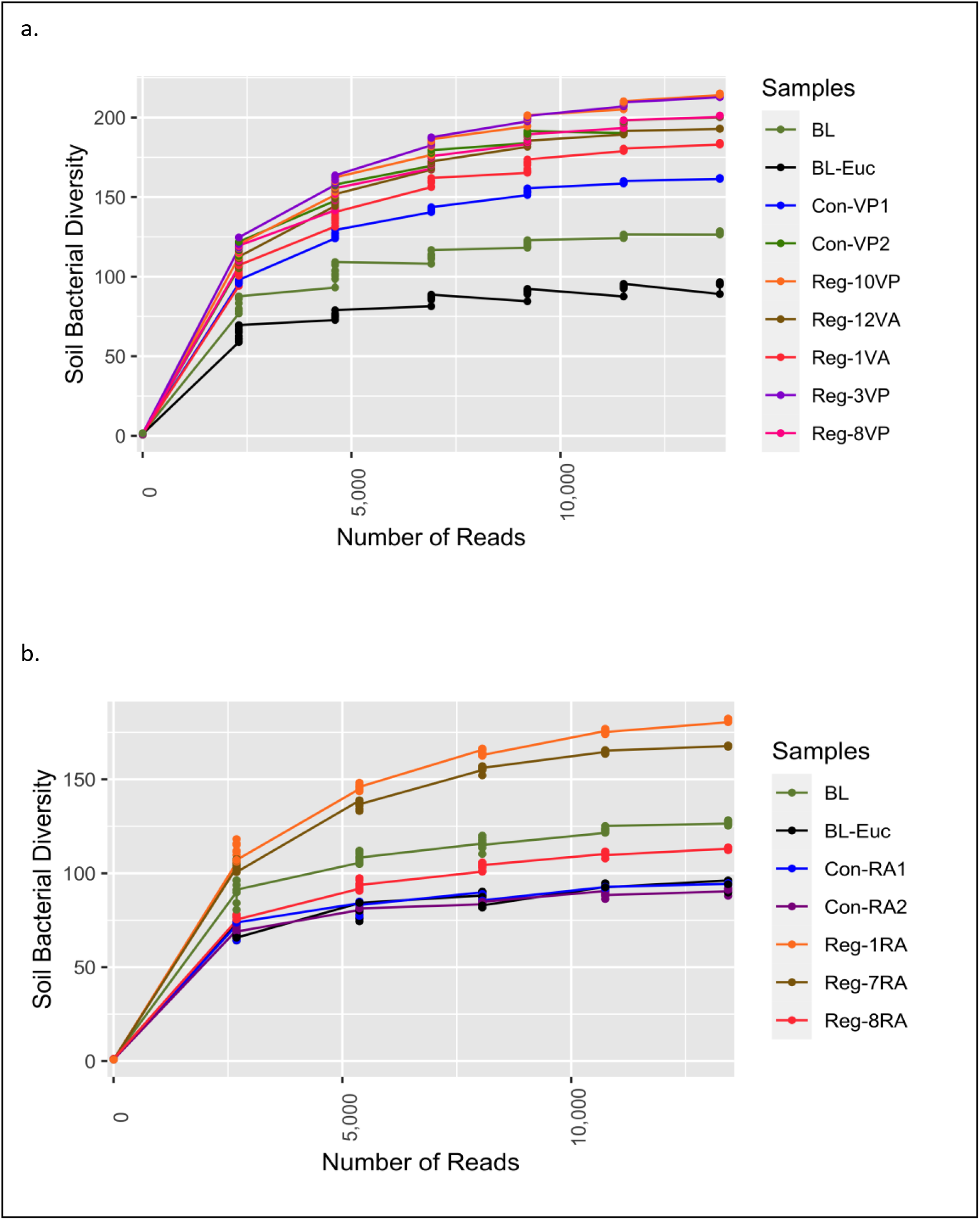
Alpha rarefaction study for soil bacterial diversity analysis of individual - **(a)** Vegetable growing Regenerative and Conventional plots with Barren land (BL) and **(b)** Ragi growing *Reg* and *Con* plots with BL.

Another interesting observation was that *Con-VP2* soil, which is exposed to a combination of conventional and regenerative practices, displayed bacterial diversity comparable to that observed in *Reg-12VA* (*Figure 2A*). This result is significant as it shows that despite merging two agricultural methods and soil sampling done in presence of crop, yet *Con-VP2* had bacterial diversity only as good as *Reg-12VA* where soil was taken in the absence of crop. Thus, a definitive augmentation in soil bacterial speciation is observed in the plots selectively practicing regenerative agriculture.

In contrast to vegetable plots, soil from the ragi growing plots could only be collected post-harvest. It is noteworthy that the CR plots displayed as poor bacterial diversity as was found in *BL-Euc* (*Figure 2b*). Least bacterial diversity in these CR plots could be due to the degradative impact of conventional fertilization on the soil’s microbial health or due to continuous cultivation with no supportive interventions or due to the inherently poor soil quality of Doddaballapur from where these soils were obtained. Interestingly, while RR plots showed better bacterial diversity than CR, the duration of regenerative practices did not correlate with the bacterial species enrichment. For example, *Reg-1RA* (practicing for 1 year) displayed higher bacterial diversity than *Reg-7RA* (practicing for 7 years). Surprisingly, *Reg-8RA* (practicing for 8 years) displayed bacterial diversity lower than even the *BL* plot. One explanation could be that at different places the starting soil will have different baselines of bacterial diversity. The sample *Reg-1RA* was collected from Magadi while *Reg-7RA* and *Reg-8RA* were obtained from Ramanagara. It seems that Magadi soil is already healthier than soil from other places owing to its mostly green-covered scape and a more recent agricultural shift in the region compared to Ramanagara, Doddaballapur, and Hosur. Therefore, soil in other places demand higher inputs to be rejuvenated compared to Magadi soil. This argument is strengthened by the finding that *Reg-3VP* (*Figure 2a*) also coming from Magadi shows a bacterial profile as rich as that observed in *Reg-10VP* plot in just three years of regenerative agriculture practice.

### Bacterial community

To elucidate the bacterial community structure in the various types of plots, we assessed and compared the bacterial phyla associated with different soil samples grouped into categories as described previously in bacterial richness and heterogeneity analysis. The major phyla observed in both kinds of vegetable plots and Barren soil included – Proteobacteria, Bacteroidota, Planctomycetota, Acidobacteriota, Chloroflexi, Actinobacteriota, Verrucomicrobiota, Cyanobacteria and Patescibacteria (*Figure 3a*). Similarly, in ragi plots and barren soil comparison the bacterial community was majorly represented by the phyla – Planctomycetota, Proteobacteria, Bacteroidota, Chloroflexi, Actinobacteriota, Acidobacteriota, Cyanobacteria, Verrucomicrobiota, Firmicutes, Patescibacteria, Myxococcota and Gemmatimonadota (*Figure 3b*). Our observations show that in regenerative agriculture plots there is a shift towards a more uniform representation of all the major phyla compared to that in conventional agriculture plots. For instance, on analysis of vegetable plots (*Figure 3a*), we see a slight reduction in the relative abundance of phyla Proteobacteriota (Barren – 16.88%, Conv – 17.45% to Reg≤3 – 16.62% and Reg>5 – 15.26%) and Acidobacteriota (Barren −9.62% Conv – 9.88% to Reg≤3 – 8.78% and Reg>5 – 7.42%) and a simultaneous increased representation of phyla – Chloroflexi (Barren – 8.75%, Conv – 6.63% to Reg≤3 – 9.11% and Reg>5 – 9.64%), Actinobacteriota (Barren – 7.02%, Conv – 6.25% to Reg≤3 – 7.80% and Reg>5 −7.15%), Cyanobacteria (Barren – 7.70%, Conv – 1.14% to Reg≤3 – 7.72% and Reg>5 – 4.47%) and Patescibacteria (Barren – 2.31%, Conv – 5.96% to Reg≤3 – 4.81% and Reg>5 – 6.73%) in regenerative soil compared to conventional and barren soil. This reorganization has led to the development of a more evenly structured community. Similarly, in the ragi plots (*Figure 3b*) we observed relatively lower levels of Acidobacteriota (Barren – 9.79%, Conv – 7.39% to Reg≤3 – 6.81% and Reg>5 – 7.02%) and higher levels of Actinobacteriota (Barren – 7.15%, Conv – 7.08% to Reg≤3 – 8.94% and Reg>5 – 14.12%) and Fermicutes (Barren – 1.89%, Conv – 2.37% to Reg≤3 – 8.01% and Reg>5 – 2.89%). Interestingly, a comparison to determine the impact of number of years of regenerative agriculture among RV plots did not show a significant change in the phylum level distribution in Reg ≤3 and Reg >5 soils. Although the comparison of RR plots (Reg >5 and Reg =1) (*Figure 3b*) showed a significantly higher representation of Firmicutes in Reg =1 soil despite only one year of regenerative practice. This is supposedly attributed to the regionally better soil of Magadi obtained Reg =1 soil (*Reg-1RA*). However, the RR plots practicing for Reg >5 years were found to show a significantly enhanced relative abundance of Actinobacteriota.

**Figure 3.**
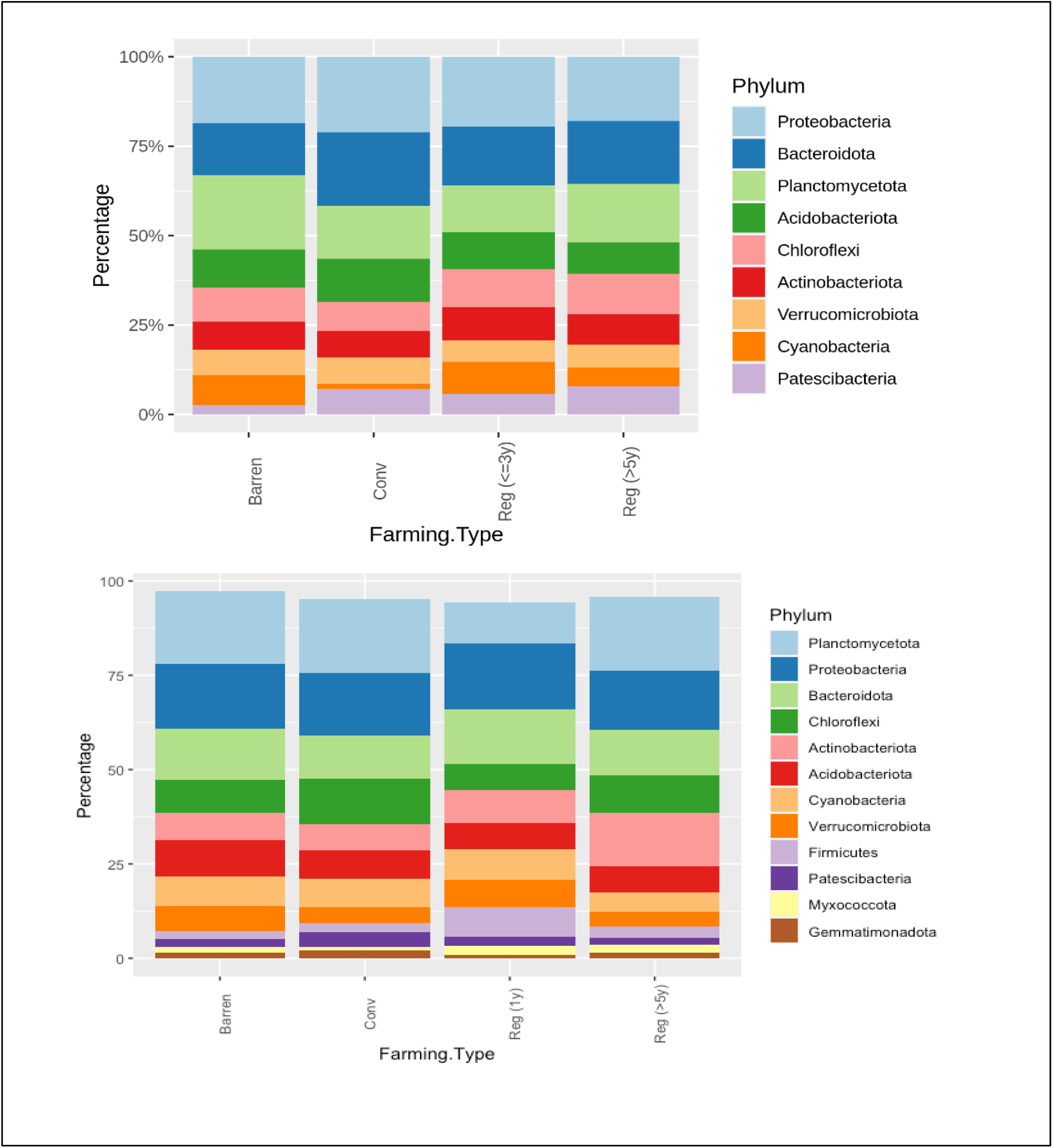
**(a)** Relative bacterial abundance at phylum levels in Conventional (*Conv*) and Regenerative (*Reg*) agriculture plots and BL in **(a)** Vegetable plots and in **(b)** Ragi plots

### PGPR community structure in regenerative agriculture

Plant Growth promoting Rhizobacteria (PGPR) are characterized to be an important group of soil bacteria that support plant growth and health by synthesizing and secreting various beneficial chemicals and nutrients in the soil. To determine the soil health in terms of PGPR representation, we selected a group of bacterial genera that have been well identified and classified as PGPRs (35, 36, 37, 38, 17). Among the genera considered here are – *Flavobacterium, Bacillus, Streptomyces, Mesorhizobium, Achromobacter, Klebsiella, Paenibacillus, Burkholderia* and *Pseudomonas*. Interestingly, RV plots when compared to CV and barren plot soils showed a relative enrichment for *Pseudomonas sp*. belonging to phylum Proteobacteria. On the contrary, RR plots demonstrated an increased representation of - *Bacillus sp*. and *Mesorhizobium sp*. The levels of *Bacillus sp*. are found to be significantly higher in both RR categories (Reg >5 and Reg = 1) compared to CR and barren land. The relative representation of *Mesorhizobium sp*. was found to be highest in Reg >5 in RR plots with a simultaneous reduction in levels of *Burkholderia sp*. compared to both CR and barren soil (*Figure 4b*). Interestingly, the genus *Streptomyces* was found to have a remarkably high representation in all Magadi plots (*Reg-1RA, Reg-1VA* and *Reg-3VP* compared to the other plots (Figure 4a, 4b). However, since we did not have any conventional plot or barren soil sample from Magadi it is impossible to estimate the contribution of RA on the enhanced *Streptomyces* configuration.

**Figure 4.**
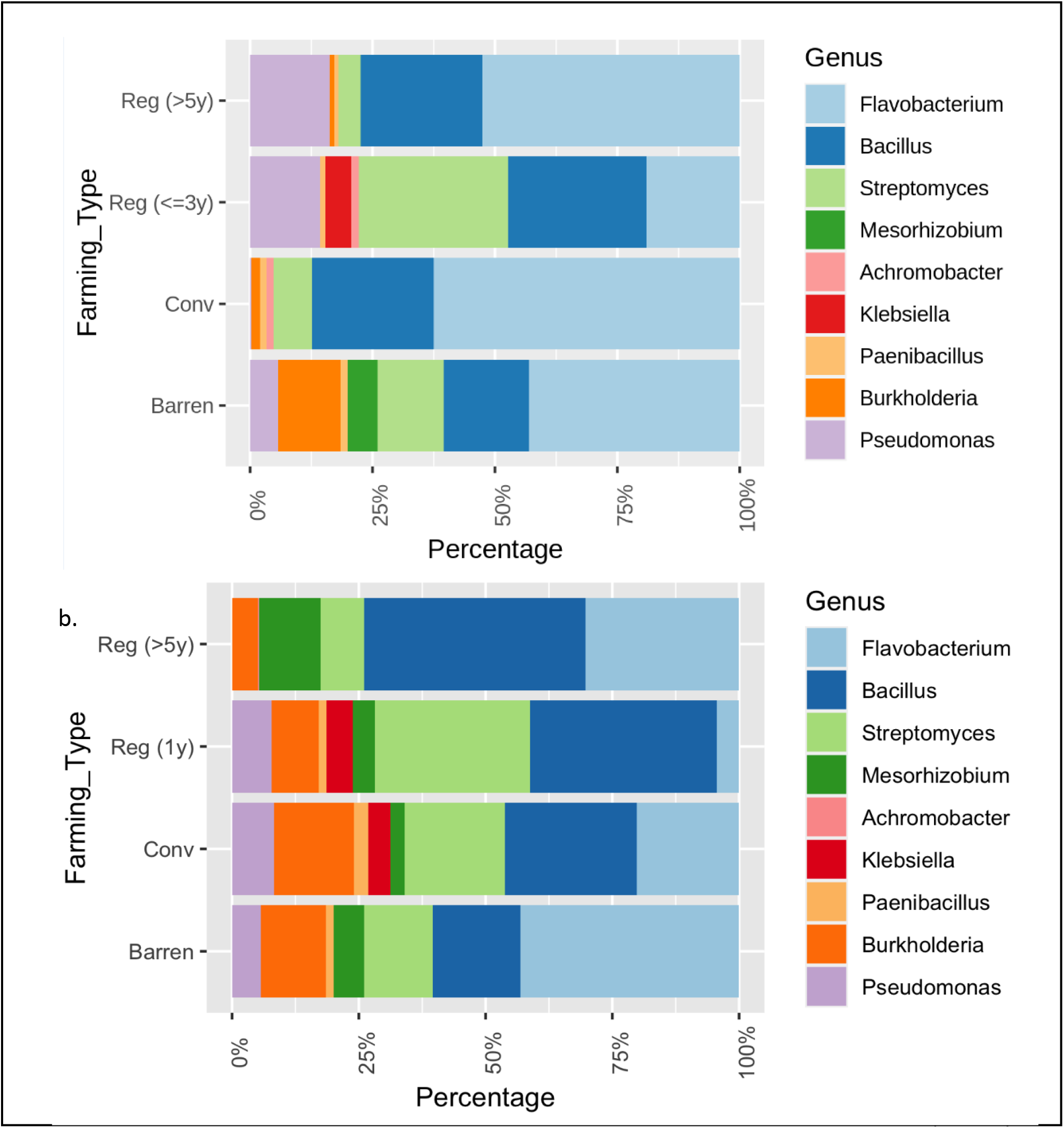
Relative composition of selected Plant Growth Promoting Rhizobacteria (PGPRs) in different soil samples. **(a)** Comparing Vegetable growing Regenerative (*Reg*) and Conventional (*Conv*) plots with BL and **(b)** Comparing ragi growing *Reg*. and *Conv*. plots with Barren.

## Discussion

Regenerative agriculture has re-emerged in the last ten years (39) as a very important means of land rejuvenation practice for sustainability in soil health, farm productivity and environmental management. Regenerative agriculture provides us with a non-synthetic, nature-based option that helps to revive the ecosystem as a whole. In India too, there is growing interest in this environmentally-safe and less expensive agriculture system, necessitating the need for elucidating its impact on soil, environment and food production as a whole. Thus, this study has attempted to decipher the impact of regenerative agriculture on soil bacterial profile, soil nutrient composition, in two cropping systems under short (<=3 years) and long-term (>5 years) influence.

### Soil Chemical Properties

Most soil samples were found to have ideal pH or a somewhat alkaline pH, which is mostly suitable for agriculture. Acidic pH was found in the soil samples coming from Doddaballapur – *BL*, *BL-Euc, Con-RA1* and *Con-RA2*. These findings are consistent with reports showing that soil from Doddaballapur generally has an acidic pH in the range from 5.0 to 7.3 (40). Highest acidity in *Con-RA1* and *Con-RA2* soils are likely due to application of synthetic fertilizers and continued cultivation without allowing the land time to revive itself (41). As per the USDA, soils with pH below 5.5 are likely to have poor calcium, magnesium and phosphorus content (32). Consistent with this, *Con-RA2* with pH<5.5 and *Con-RA1* exhibiting pH around 5.5 showed low levels of calcium, magnesium and phosphorus. We further observed that soil samples with pH values above 7.8 have adequate calcium and magnesium levels but depleted copper, manganese and iron content. This was found to be somewhat true for the samples – *Reg-10VP* (pH = 7.95) and *Reg-8VP* (pH = 8.31) where calcium and magnesium levels are in surplus, whereas copper is much above the ideal limit of 0.2 ppm. Most regenerative agriculture plots were found to have ideal or slightly alkaline pH levels.

High phosphorus levels in conventional agriculture plots (*Con-VP1* and *Con-VP2*) is most likely attributed to the excessive chemical - NPK fertilization where phosphorus and potassium remain in the soil over time whereas nitrogen gets lost due to leaching and nitrogen cycling (42, 43, 44). Available literature shows that as soil degrades there is a simultaneous decline in the composition of all its nutrients (45). However, since the BL soils considered in this study did not show a marked reduction in any of the nutrients, therefore these soils may not be suitably classified as degraded. Although, it may be interesting to study the microbial health and nutrient composition of these soils in a span of 3-5 years from now, to observe the changes in the barren soil composition to estimate the progression of degradation.

### Bacterial richness and diversity

As shown by multiple studies from across the world, we found that regenerative agricultural system improves bacterial diversity compared to both conventional and barren soil (47, 49, 50, 55, 56, 14). Here we report an increase in bacterial richness and heterogeneity across all regenerative plots, including those that have moved to this system very recently. This is a very significant result indicating that application of regenerative agriculture, from the outset boosts and modulates the soil’s bacterial growth, promoting a more heterogeneous composition for carrying out various soil health enhancing activities. Another important finding from the alpha diversity comparison of vegetable plots is that longer the period of RA application greater is the community’s bacterial diversity. These findings confirm the biological enrichment abilities of regenerative agriculture (6, 10).

The demonstrated lower alpha diversity among RA plots with no crops during soil sampling versus those with crops underpins the fact that roots of the crops induce proliferation of a large variety of root colonizing and plant growth stimulating rhizosphere microbes (33, 34). Although the RR plots also showed the highest alpha diversity compared to CR and BL, yet a reverse time-dependence trend was observed among the ragi RA plots. This could be attributed to the soil sampling done in the absence of crop and the regional differences contributing to a disproportional decline in microbial community profiles. In addition, inherent regional soil characteristics and composition may also play a significant role in shaping the microbial community structure (50). This is evident from the Magadi obtained soils - *Reg-3VP* and *Reg-1RA*, which displayed highest alpha diversity in their respective groups (*Figure 2a, 2b*).

Among all the RA plots in this study *Reg-10VP* was observed to show the best overall profile in terms of both bacterial community structure as well as soil physicochemical characteristics. Looking at the nutrient and bacterial profile of sample *Reg-10VP*, one can construe that continued regenerative practice over five years or more has the capability to improve the soil’s bacterial community structure, which would in turn enhance soil and plant health. We know from the farmer interviews that *Reg-10VP* has been demonstrating good crop yield. Furthermore, it is interesting to note that the Potassium, Phosphorus and Soil Organic Carbon (SOC) content of this soil is better than that of other farms. Studies have claimed that regenerative agriculture is the most promising way to sequester atmospheric carbon and mitigate climate change (51, 52, 53). India’s soil is reported to be highly depleted in SOC levels (54). A time series comparison of organic agriculture with conventional has shown that organic practice has helped improve SOC levels in soil from 12.5 g/dm^3^ to 21 g/dm^3^ and microbial biomass from 87 mg/kg to 120 mg/kg in a span of just one year (57). An all-round improvement in soil bacterial and nutrient profile displayed by *Reg-10VP* holds a similar promise for regenerative agriculture in India. The carbon enriched *Reg-10VP* soil confirms the potential of regenerative agriculture in boosting carbon sequestration. Going by this argument, Indian agricultural land can form one of the largest terrestrial carbon sinks to reverse climate change. These findings suggest that regenerative practices stimulate the formation of a healthy microbial community with diverse species to carry out the biogeochemical processes more efficiently, providing a buffering mechanism that overcomes the pressures of the ecosystem. These resilient ecosystems can easily tackle the vulnerabilities due to nutrient inadequacies, pathogen and pest attacks as well as climate change (14).

The intermediate level of bacterial diversity in CV plots is most likely due to the mixed agriculture methods used by these farmers. Here the farmers integrate both organic manure and chemical fertilization methods to accrue the benefits from both the systems. If used judiciously, the synthetic fertilizers may also be useful to supplement the soil with necessary nutrients and in maintaining the soil’s organic matter (SOM) (9, 12, 41). BL soil’s poor bacterial richness and high evenness is attributed to absence of any vegetation for multiple years resulting in continued exposure to weathering, erosion and deterioration (58). Thus, the BL soil has become depleted in its microflora and enriched in fewer robust microbes that can sustain in harsh conditions. Studies conducted on degraded soil in China reveal that poor quality soils display a depleted Operational Taxonomic Unit (OTU) richness for beneficial microbes and significant enhancement of pathogenic microbes (59).

### Bacterial community structure

In RV plots we observed an increased representation of Chloroflexi, Cyanobacteria, Patescibacteria and a slight increment in Actinobacteriota. Enrichment for Cyanobacteria generally will have a beneficial impact on soil health as these bacteria improve soil fertility by fixing nitrogen, phosphorus and carbon and by producing plant growth promoting hormones and siderophores (60). Additionally, exopolysaccharides, which form 25% of the total biomass of Cyanobacteria are capable of aggregating the soil and organic content and improving the soil’s water retention capacity (61, 62). Thus, Cyanobacteria improve the soil’s physical and chemical properties, promoting plant growth and productivity. Cyanobacterial bio-fertilizer comprising a mixture of free-living Cyanobacteria is highly recommended for biological nitrogen fixation and phosphorus mobilization in rice and wheat fields, contributing to significant increase in plant biomass, grain yield and nutritive value (61). Patescibacteria and Actinobacteriota have been suggested to induce plant root biomass and thus supporting better nutrient acquisition (63). Role of Chloroflexi in plant health is not clear although study has reported that Chloroflexi comprising anaerobic bacteria, are found to be enriched in paddy fields depending on oxygen availability and regulate soil bacterial community composition (64).

Likewise, the RR plots showed an enrichment for Firmicutes and Actinobacteriota population, which again form a group of extremely beneficial plant growth promoting bacteria (65). Phylum Firmicutes comprises a number of agro-ecologically beneficial bacterial genera, such as *Bacillus, Paenibacillus*, *Lysinibacillus*, *Brevibacillus*, *Planococcus*, *Clostridium*, *Sporosarcina* etc. (65). Many of these bacterial genera (eg. Bacillus) have been identified as biocontrol and phytoremediation agents and others as Plant Growth Promoting Rhizobacteria (PGPRs). Thus, enrichment for Firmicutes in regenerative agriculture plots signifies a marked improvement in soil health. Members of the phylum Actinobacteriota like *Streptomyces, Brevibacteria* and *Nocardia* promote plant growth as bio-fertilizers and bio-controllers for agricultural sustainability (66). Similarly, a study has also shown the significance of both Firmicutes and Actinobacteriota in controlling bacterial disease incidence in tomato plant (67).

Barren soil was observed to have a relatively higher representation of Planctomycetota compared to both conventional and regenerative soils. Additionally, we observed a higher level of phylum Acidobacteriota representation in barren soil when compared with CR and RR plots. This is in coherence with a report where an increase in relative abundance of Proteobacteria, Acidobacteriota and Bacteroidota was observed in degraded soils whereas healthy soils were enriched for Actinobacteriota and Firmicutes (45).

### Plant Growth Promoting Rhizobacteria (PGPRs)

New developments in the field have shown that healthy soils are enriched in Plant Growth Promoting Rhizobacteria (PGPRs). These PGPRs secrete plant growth hormones and regulatory chemicals in the rhizosphere, facilitating plant growth by enabling plant nutrient procurement, modulating plant hormone levels and by releasing biocontrol agents to protect plants against pathogens. Many bacterial genera including *Pseudomonas, Bacillus, Streptomyces, Flavobacterium, Achromobacter, Mesorhizobium, Paenibacillus, Sinorhizobium, Burkholderia, Rhizobium*, etc. have been classified as PGPRs. Many of these bacteria are being currently used as biocontrol agents and as bio-fertilizers (38, 68, 69, 70, 71, 72, 73). Augmentation of these bacterial genera in soil directly indicate towards improvement in soil health.

Our study showed a relative enrichment for *Pseudomonas sp*., in RV plots, *Bacillus sp*., and *Mesorhizobium sp*. in RR plots. Many studies have provided evidence that *Pseudomonas* forms the core of PGPRs for many vegetable, fruit and flowering plants (72, 73). According to studies, *Pseudomonas* is the most efficient producer of ammonia and enhances bioavailability and bioassimilation of nutrients, promoting plant growth and yield (73). Thus, enrichment for *Pseudomonas sp*. is essentially a favorable development in RV plots. Interestingly, studies show that ragi growth is promoted by the rhizospheric growth of *Bacillus sp*. The *Bacillus sp*. support ragi growth by fixing nitrogen and protecting the crop against the foot-rot disease causing pathogen, *Sclerotium rolfsii* (74). Furthermore, *Bacillus sp*. are known to be involved in improving the nutritive value of the ragi grains by enriching them with essential amino acids (75). An Ethiopian study suggests that *Bacillus* and *Pseudomonas* species form significant PGPRs supporting vegetable crops (72). In effect an enrichment for *Pseudomonas sp*. in RV plots and for *Bacillus sp*. in ragi plots signify a beneficial transformation in soil bacterial composition. Likewise, *Mesorhizobium sp*. are found to be very useful PGPRs with their special property of synthesizing ACC deaminase enzyme which protects plant against abiotic stress by degrading ACC which forms the precursor for ethylene. Additionally, *Mesorhizobium sp*. synthesizes IAA which promotes plant root growth and also is involved in inorganic phosphate solubilization making it available to plants (76). Thus enrichment for *Mesorhizobium sp*. has multifarious benefits. Magadi soil seems to be inherently enriched in *Streptomyces sp. Streptomyces sp*. also form an important group of agriculturally beneficial rhizosphere bacteria (77, 78). *Streptomyces* synthesize plant hormone – Indole acetic acid (IAA) in moderate quantities and help in phosphate solubilization and stress tolerance thus boosting plant growth and productivity. Thus this clearly indicates that regenerative agriculture practices are able to induce a healthy microbial population in the soil for promoting soil’s overall health and agricultural.

### Regenerative practices and their impact

Almost all regenerative agricultural plots considered here have indicated to the use of farmyard manure as an important supplement for soil management. Manure addition has been ascribed to inducing increased microbial biomass in soil (79, 80). Some studies indicate that the type and source of farm manure dictates the soil microbial population (59, 60). However, it may be difficult to define the source of origin of a microbe in soil. For instance, one report claims that cow manure enriches the soil for Firmicutes and Bacteroidota while another suggests an enrichment for Firmicutes and Proteobacteria. Contrary to this, a recent study claims that in a span of two weeks from manure addition, the microbes coming from the manure are mostly lost while the soil-borne microbes are activated to grow and multiply (81). Regenerative plots demonstrated an increased growth of Firmicutes particularly *Bacillus sp*. in ragi fields and Proteobacteria (*Pseudomonas sp*.) in vegetable plots. In addition, since almost all the regenerative farms are using multiple regenerative practices apart from just farmyard manure application, these additional treatments will also influence the soil microbiome. More studies are therefore required to ascertain the roles of these individual treatments in determining the microbial community structure. In *Reg-10VP* plot a rich supplementation of farmyard manure (400 kg/row) could have been a significant contributing factor to the plot’s best nutrient and bacterial profile. However, since not all farms will be able to afford this kind of soil supplementation regimes, policies and practices such as encouragement of circular economy to provide household based compost to farmers is necessary.

### Influence of region and crop on soil bacterial composition

Soil microbial community structure was found to be influenced by regional and spatial characteristics. Certain regions required greater inputs with many years of application and others much less to achieve a credible improvement in microbial health and soil quality. This is evident from the Magadi obtained soil samples – *Reg-1VA, Reg-3VP* and *Reg-1RA*. These regenerative agriculture plots have been practicing for just one, three and one year respectively, yet these soils showed very high alpha diversity (*Figure 2a, 2b*) and a distinctly heterogeneous and highly diverse bacterial composition with a higher representation of *Streptomyces* sp. (*Figure 4a, 4b*). Additionally, crop-plants also play a role in defining the soil’s bacterial community structure as is evident from the varied profiles exhibited by regenerative plots growing ragi and vegetable crops (82, 83, 84).

### Soil Microbiome Impacts of merging conventional and regenerative systems

The *Con-VP2* where soil sampling was done in presence of crop forms a suitable example of a plot where the two agricultural systems – Conventional and Regenerative have been integrated for land and crop management. This plot shows a distinctly high alpha diversity comparable to that in *Reg-12VA* plot, where soil was collected in absence of crop. However, the alpha diversity of *Con-VP2* is still found to be lesser than all the RV plots where soil was taken in the presence of the crop. Thus we conclude that addition of any amount of synthetic fertilizer will have an adverse impact on the soil microbiome. Application of inorganic fertilizers comes with a host of adverse effects in soil including increase in salinity, acidification, soil compacting and poor water retention, impact on biogeochemical processes by altering microbial dynamics, accumulation of toxic wastes/heavy metals and finally reduced microbial diversity (85). Ragi conventional plots obtained in our study are a clear indication of the detrimental effect of conventional agriculture. In this study, merging of the two systems of agriculture shows an intermediate profile in terms of bacterial diversity, however based on available literature it would be safer to adopt regenerative agriculture independently for sustainability.

## Conclusion

This study aimed to compare and elucidate the effectiveness of regenerative agriculture practice on soil microbial and nutritive health with respect to conventional agriculture and barren soil. Barring a few exceptions owed to different original baselines of the selected plots, the observations show that extended periods of regenerative practice does improve soil bacterial diversity and soil nutrient health. Even SOC levels were found to be within the desired range in long-term regenerative application plots. Regenerative plots showed an enrichment for bacterial phyla which promote soil health and plant growth sustainably. Despite variabilities in regenerative practices adopted by the farmers we could still see a better bacterial community structure and richness in all regenerative plots. The results reinforce the importance of regenerative agriculture for sustainable management of soil health and agriculture. Thus we conclude that at least five years and more of regenerative agriculture practice can help to boost soil microbial health potentiating an enrichment for major and micronutrients, subsequently enhancing plant growth and productivity. Furthermore, we conclude that mixing of the conventional and regenerative practices is not a sustainable option for maintaining good biological health of the soil.

The RA plot showing the best bacterial profile and ideal SOC levels uses very heavy application of farmyard manure for soil management and Jeevamrutha for pest management. Thus although regenerative agriculture has the ability to induce beneficial outcomes in soil health and agriculture, the required impact is made possible only with a heavy use of amendments at least in the initial decade or so. This identifies the need for instituting a continued and surplus supply of manure to the farmers for ensuring high grade outputs.

## Acknowledgements

This research is part of the SHEFS - an interdisciplinary research partnership, forming part of the Wellcome Trust’s funded Our Planet, Our Health programme, with the overall objective to provide novel evidence to define future food systems policies to deliver nutritious and healthy foods in an environmentally sustainable and socially equitable manner. This research was funded by the Wellcome Trust through the Sustainable and Healthy Food Systems (SHEFS) Project (Grant number-205200/Z/16/Z).

## Notes

### Competing Interest Statement

The authors have declared no competing interest.

